# Hidden hematological, biochemical and immune costs of asymptomatic malaria infections in semi-wild chimpanzees

**DOI:** 10.1101/2025.11.04.686503

**Authors:** A. Nowakowski, E. Elguero, C. Patterson, A. Boissière, F. Degrugillier, C. Sidobre, C. Arnathau, P. Grentzinger, E. Willaume, A. M. Talman, B. Malleret, L. Boundenga, B. Ngoubangoye, F. Prugnolle, S. C. Wassmer, V. Rougeron

**Affiliations:** REHABS, International Research Laboratory, CNRS-NMU, George Campus, Nelson Mandela University, Madiba Drive, 6529 George, South Africa; Laboratory MIVEGEC, University of Montpellier, CNRS, IRD, 900, rue Jean-François Breton, 34900 Montpellier, France; Department of Infection Biology, London School of Hygiene & Tropical Medicine, London, UK; Laboratory Animal, Santé, Territoire, Risque, Ecosystème (ASTRE), UMR 117, Campus International de Baillarguet, CIRAD, Montpellier, France; Sodepal, La Lékédi Park, Bakoumba, Gabon; Department of Microbiology and Immunology, Immunology Translational Research Programme, Yong Loo Lin School of Medicine, Immunology Programme, Life Sciences Institute, National University of Singapore, 117597 Singapore, Singapore; Singapore Immunology Network (SIgN), Agency for Science & Technology, Singapore, Singapore; CIRMF, Centre Interdisciplinaire de Recherches Médicales de Franceville, Franceville, Gabon; Department of Anthropology, Durham University, South Road, Durham, DH1 3LE, UK; Sustainability Research Unit, George Campus, Nelson Mandela University, Madiba Drive, 6529 George, South Africa

## Abstract

The health consequences of *Plasmodium* infections in wild great apes, particularly in asymptomatic animals, remain poorly understood. This study investigated the hematological and immune impacts of natural malaria infections in 27 semi-wild chimpanzees (*Pan troglodytes troglodytes*) from Gabon. Using MinION sequencing and species-specific PCR, results showed a 48.15% overall *Plasmodium* infection rate, with frequent multi-species co-infections involving *Plasmodium gaboni*, *Plasmodium reichenowi*, and *Plasmodium vivax-like* parasites. In addition, younger animals were significantly more infected and exhibited higher parasitemia levels, especially those with triple infections involving *P. vivax-like*. Profiling of 15 hematological markers and 8 cytokines/chemokines known to be associated with malarial infections in humans revealed significant alterations in infected chimpanzees, including elevated urea, reduced creatinine, and increased systemic concentrations of pro-inflammatory (TNF, IL-1β, CCL3) and anti-inflammatory (IL-10) cytokines. *Ex vivo* PBMC stimulation yielded higher IL-10 in infected than non-infected individuals, indicating a regulatory-skewed cytokine response at the time of sampling. These results suggest that malaria in chimpanzees is associated with systemic immune modulation and accompanied by signs of physiological stress, including potential renal dysfunction. This study challenges the assumption that chronic *Plasmodium* infections are entirely benign in great apes and highlight the need to integrate immunological health indicators into conservation strategies. Broader immune profiling and longitudinal studies will be essential in the future to assess long-term health outcomes and resilience in these endangered populations.

## Introduction

Malaria is a vector-borne disease caused by protozoan parasites of the genus *Plasmodium* [1]. Six species are currently recognized as infecting humans: *Plasmodium falciparum*, *Plasmodium vivax*, *Plasmodium malariae*, *Plasmodium ovale wallikeri*, *Plasmodium ovale curtisi*, and *Plasmodium knowlesi*. African great apes are similarly natural hosts to several *Plasmodium* species closely related to those infecting humans. These parasites are divided into two subgenera: (i) *Plasmodium*, which includes all human-infecting species except *P. falciparum*, and three species infecting apes, *Plasmodium vivax*-like, *Plasmodium malariae*-like, and *Plasmodium ovale*-like; and (ii) *Laverania*, which includes *P. falciparum* and six ape-specific species, *Plasmodium praefalciparum*, *Plasmodium reichenowi*, *Plasmodium billcollinsi*, *Plasmodium blacklocki*, *Plasmodium adleri*, and *Plasmodium gaboni*, that infect chimpanzees, gorillas, and/or bonobos [2–4].

While the health impacts of *P. falciparum* and *P. vivax* infections in humans are well documented, the consequences of *Plasmodium* infections in great apes currently remain poorly understood. This knowledge gap largely stems from the logistical challenges of monitoring natural infections in non-human primates (NHPs), including difficulties in sampling protected species and limited access to field-based laboratory facilities. Although some studies suggest that these infections are often asymptomatic in great apes, others have reported malaria-related clinical signs in chimpanzees, indicating potential variability in disease outcomes [3,5–12]. Historically, the first documented descriptions of malaria parasite infections in African great apes date back to the 1910-20s [13–15]. Rodhain’s studies reported a lack of illness signs in chimpanzees (*Pan troglodytes troglodytes*) infected with either great ape or human *Plasmodium* strains [13]. Conversely, one study described a juvenile chimpanzee infected with *P. falciparum* and *P. vivax* that had fever and loss of appetite [14]. Despite limited and sometimes conflicting evidence, the prevailing view was not that chimpanzees fail to get infected, but that naturally acquired *Plasmodium* infections in chimpanzees are usually subclinical, with overt malaria illness considered rare in the wild. It was only decades later that sporadic reports began to document *Plasmodium* infections in chimpanzees housed in sanctuaries or captive settings, though these accounts also yielded inconsistent conclusions. For example, Hayakawa et al. (2008, 2009) reported chronic *P. malariae*-like infections in two captive chimpanzees in Japan who remained asymptomatic for over 30 years [16,17]. Similarly, Krief et al. (2010) reported *P. falciparum* infections in bonobos from the Democratic Republic of Congo without any clinical signs [18]. Conversely, other studies have documented symptoms in African great apes, such as Tarello (2005), who reported the death of a one-year-old chimpanzee infected with *P. reichenowi*, although the exact cause of death remained unclear [19].

To date, the only study to our knowledge documenting clinical signs associated with *Plasmodium* infection in African great apes is that of Herbert et al. [12], which reported anemia and hyperthermia in a juvenile chimpanzee naturally infected with *P. reichenowi* in Gabon. This case suggested that, under certain conditions, *Plasmodium* infections may have measurable health impacts (i.e. fever and strong anemia). However, it remains the sole documented instance of apparent malaria-associated pathology in chimpanzees, and no subsequent studies have definitively addressed the clinical consequences of natural infections in wild great apes. As a result, the research community remains divided (among others, [3,5,20]). As the paucity of systematic clinical monitoring, current evidence cannot resolve the full spectrum of outcomes. Rather than a simple “severe vs. innocuous” dichotomy, effects likely vary with age and exposure (acquired immunity), parasite lineage and parasitemia, host species/individual condition, and co-infections/nutritional stress, so mild or subclinical impacts may be common yet under-detected in cross-sectional studies.

Given the ongoing debate surrounding the health impacts of malaria in African great apes, this study aimed to assess the hematological and immune consequences of natural *Plasmodium* infections in chimpanzees in Gabon. In humans, various hematological parameters are well-established indicators of malaria severity and parasite virulence during the asexual blood stage. Infections are typically associated with reduced platelet, leukocyte, lymphocyte, red blood cell, hematocrit and hemoglobin counts, as well as elevated monocyte and neutrophil counts [21]. Because malaria also frequently affects the kidneys and liver in humans, renal (urea, creatinine) and hepatic (ALAT, ASAT, GGT) biochemical markers, along with lipid metabolism markers (cholesterol, triglycerides), were included as key indicators of organ function and metabolic status [22,23]. Additionally, immune responses to *P. falciparum* in humans are characterized by the secretion of both pro- and anti-inflammatory cytokines to respectively control parasitemia and modulate immunopathology [24]. Based on these observations, we selected a panel of 15 hematological biomarkers (described above), as well as eight cytokine and chemokine markers selected based on their known role in malarial infections in humans (TNF, IL-6, IFN-γ, IL-1β, IL-4, CCL3/MIP-1α, CCL5/RANTES, and IL-10), for analysis in infected and non-infected chimpanzees. This comprehensive dataset enables the first detailed assessment of the hematological, biochemical and immunological effects of *Plasmodium* infection in semi-wild chimpanzees, providing meaningful comparisons with human malaria for the first time.

## Results

### *Plasmodium* species identification from MinION reads and specific *P. vivax-like* PCR

In this study, we screened 27 chimpanzee samples for *Plasmodium* infection using amplified *Plasmodium* cytochrome-b (cyt-b) gene coupled with MinION nanopore sequencing strategy to determine the infection status and species composition of individual infections. For each sequenced sample, a random subset of 10,000 reads was analyzed, and a threshold of 300 reads (3%) was applied to distinguish true positives from background noise. This threshold accounts for the relatively high error rates associated with nanopore sequencing (10–15%), which aligns with prior studies in clinical diagnostics and metagenomics that use similar cutoffs to filter out low-confidence detections [21,22]. Based on this threshold, all *Plasmodium*-positive identifications were of high-confidence and supported by a robust read count. Results revealed that co-infections involving *P. gaboni* (PG) and *P. reichenowi* (PR) were frequent, and only one animal had a mono-infection with a *P. ovale*-like (PO) parasite (Supplementary Table 1). To overcome the limited sensitivity of cyt-b for detecting *P. vivax-like* (PV) parasites, we additionally performed a PCR targeting the mitochondrial cytochrome oxidase I (cox1) gene (Liu et al.) [25,26], and combined these results to estimate an overall *Plasmodium* infection prevalence of 48.15% (13/27; all species pooled; Figure 1A). Among the infected chimpanzees, 25.91% (7/27) had tri-infections with *P. gaboni*, *P. reichenowi*, and *P. vivax*-like (PG/PR/PV), 18.54% (5/27) had bi-infections with *P. gaboni* and *P. reichenowi* (PG/PR), and 3.7% (1/27) had a mono-infection with *P. ovale*-like (Figure 1A).

**Figure 1.**
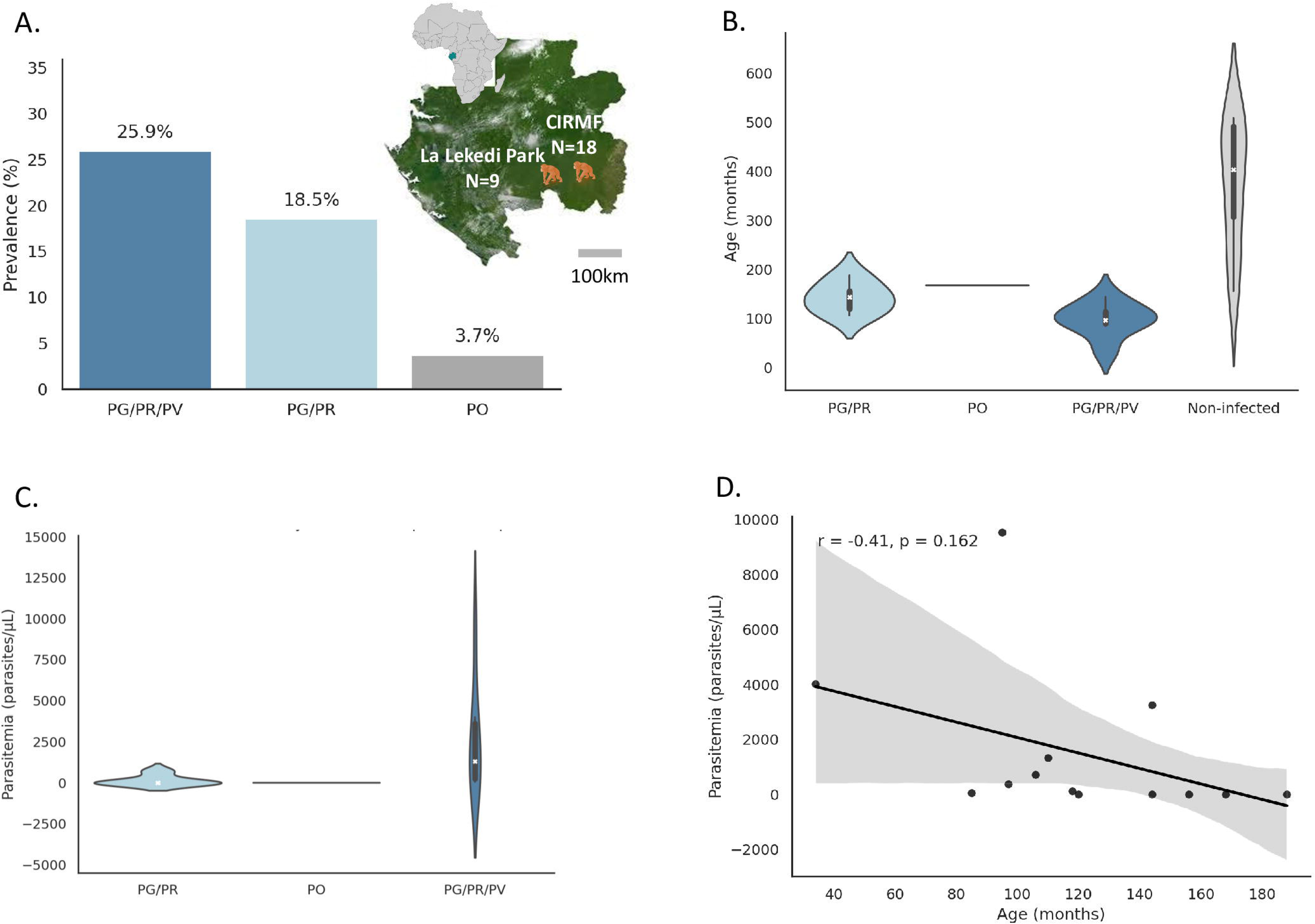
Geographic origin, *Plasmodium* prevalence, age distribution, and parasitemia levels in 27 *Pan troglodytes troglodytes* from Gabon. (A) Geographic origin of sampled chimpanzees. Animals were sampled from two sites in Gabon: the Centre Interdisciplinaire de Recherches Médicales de Franceville (CIRMF; *N* = 18) and La Lékédi Park (*N* = 9). The adjacent bar plot shows *Plasmodium* infection prevalence by species group: PG/PR/PV (*P. gaboni*, *P. reichenowi*, *P. vivax*-like), PG/PR (*P. gaboni*, *P. reichenowi*), and PO (*P. ovale*-like). (B) Age distribution (in months) by infection status: non-infected (light grey), PG/PR-infected (light blue), PG/PR/PV-infected (steel blue), and PO-infected (dark grey). Violin plots display full distributions with internal boxplots showing medians and interquartile ranges; the PO-infected animal is represented by a single point (*n* = 1). (C) Parasitemia levels (parasites/μL of blood) by infection group. Violin plots show the distribution of parasitemia in PG/PR-, PO-, and PG/PR/PV-infected chimpanzees, with greater variability observed in the PG/PR/PV group. (D) Correlation between age and parasitemia in infected chimpanzees. Each point represents one animal. A moderate negative correlation is observed (r = –0.41, *p* = 0.162), suggesting younger animals tend to have higher parasitemia. The regression line is shown in black with a shaded 95% confidence interval.

### Age impact on *Plasmodium* prevalence and multi-species infections consequences on parasitemia in chimpanzees

We analyzed the influence of several host factors, such as age, sex, body temperature, and blood group, on the *Plasmodium* infection status and parasitemia levels in chimpanzees. While body temperature, sex, and blood group showed no significant differences between infected and non-infected animals, age emerged as a key factor associated with infection (Supplementary Table 2). Infected chimpanzees were significantly younger than non-infected ones (median age: 118 months (9.8 years) vs. 380 months (31.7 years); t-test *P* < 0.05; Figure 1B; Supplementary Table 2). This pattern was particularly pronounced in animals infected with *P. gaboni* and *P. reichenowi* (PG/PR), and with all three species (*P. gaboni*, *P. reichenowi*, and *P. vivax*-like; PG/PR/PV), who were significantly younger than non-infected chimpanzees (Figure 1B). Analysis of parasitemia levels by qPCR further revealed that multi-species infections were associated with higher parasite burdens. Parasitemia ranged from as low as 5.11 copies/μL in a mono-infected *P. ovale*-like animal to a median of 7.79 copies/μL in bi-infected chimpanzees (PG/PR) and a substantially higher median of 1323.90 copies/μL in tri-infected animals (PG/PR/PV) (Figure 1C). This difference was statistically significant (Mann-Whitney U = 3; *P* < 0.05). Moreover, parasitemia tended to be higher in younger animals, as shown by regression analysis (Figure 1D).

### Blood and immunological parameters associated with *Plasmodium* infection status

To assess global variation in hematological, biochemical and immune responses to *Plasmodium* infection, a principal component analysis (PCA) was performed on 15 hematological and 8 cytokine and chemokine parameters. When comparing all infected chimpanzees (PG/PR/PV, PG/PR and PO) to non-infected animals, blood parameters PCA showed partial separation along PC1 and PC2 (21.88% and 16.84% variance, respectively; Figure 2A). In contrast, cytokine/chemokine-based PCA showed a more pronounced discrimination (PC1 = 50.36%, PC2 = 20.34%; Figure 2B). Stratification by infection group (PG/PR, PG/PR/PV, PO) emphasized some less clear species-specific patterns. While blood parameters PCA did not show clear clustering between non-infected, PG/PR-, and PG/PR/PV-infected animals (Figure 2C), cytokine/ chemokine parameters PCA revealed some separation of PG/PR infected animals from non-infected ones along PC2 (20.34% variance; Figure 2D).

**Figure 2.**
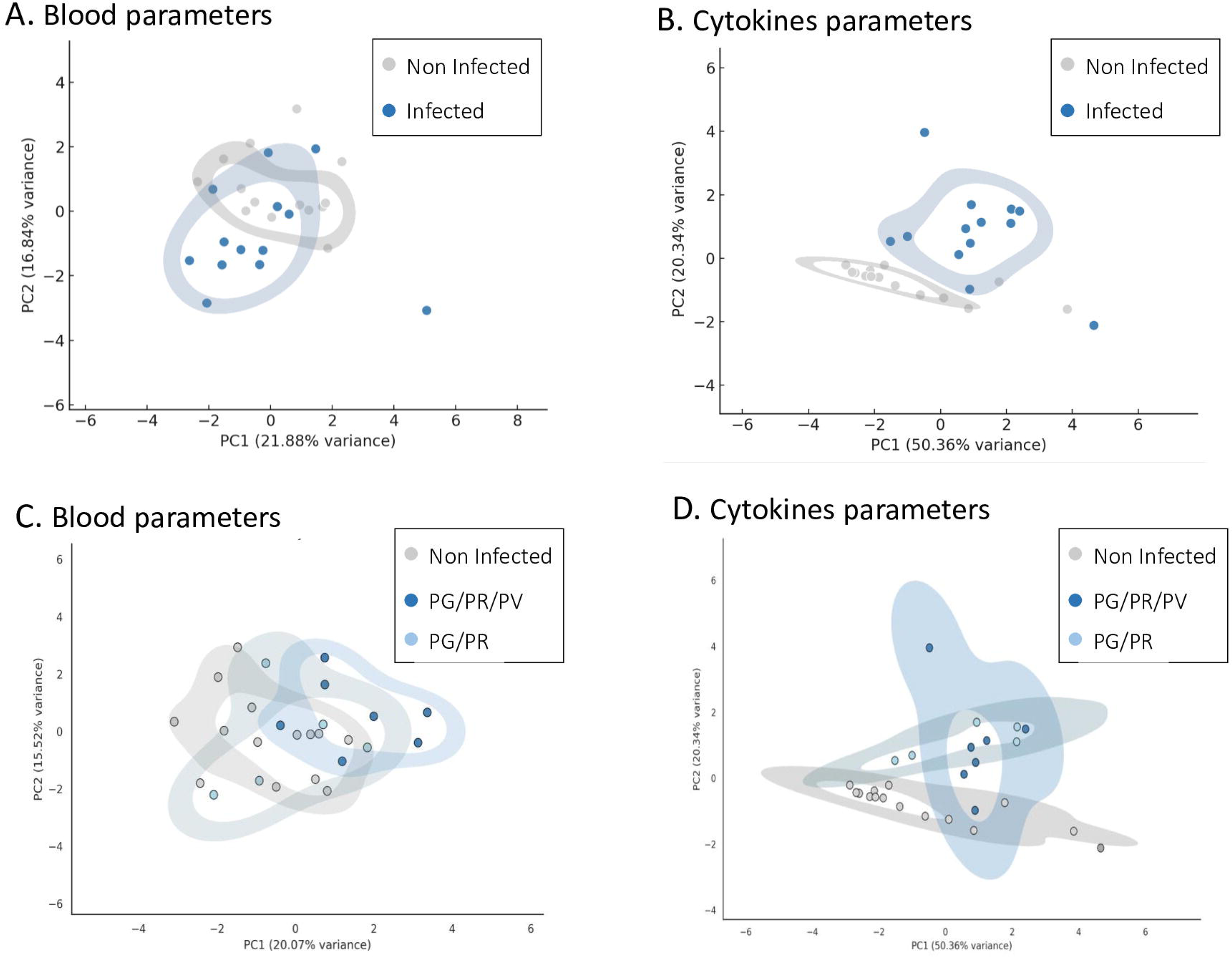
Principal Component Analysis (PCA) of blood and cytokine parameters by *Plasmodium* infection status and species in chimpanzees. (A) PCA of 15 hematological and biochemical parameters comparing non-infected and *Plasmodium*-infected chimpanzees. (B) PCA of 8 cytokine and chemokine parameters comparing non-infected and infected chimpanzees. (C) PCA of hematological and biochemical parameters stratified by infection group: non-infected, PG/PR-infected (*P. gaboni*, *P. reichenowi*), and PG/PR/PV-infected (*P. gaboni*, *P. reichenowi*, *P. vivax*-like). D. PCA of cytokine and chemokine parameters stratified by the same infection groups. In all panels (A–D), prediction ellipses represent each group at an 80% confidence level, illustrating the separation between infection categories.

Univariate analyses identified six parameters significantly associated with infection. Infected chimpanzees had higher urea (P-value = 0.012; mean = 0.80 mmol/L, vs 2.17 mmol/L) and lower creatinine (P-value = 0.014; mean = 86.11 μmol/L, vs 67.63 μmol/L) compared to non-infected chimpanzees (mean = 2.17 mmol/L and 67.63 μmol/L, respectively) (Table 1, Figure 3A, Supplementary Figure 1). In addition, the urea-to-creatinine ratio was higher in infected chimpanzees (ratio = 0.03) compared to non-infected individuals (ratio = 0.009). Cytokine/chemokines markers IL-10 (P-value = 0.002), CCL3 (P-value = 0.001), TNF (P-value = 0.001), and IL-1β (P-value = 0.003) were also elevated in infected animals (Table 2; Figure 3B; Supplementary Figure 2). Upon stratification by infection group (PG/PR, PG/PR/PV), the same six parameters remained significant, and in addition white blood cells (WBC) count also became significantly associated with infection status (P-value = 0.045), with lower counts in PG/PR infected chimpanzees but higher in PG/PR/PV infected chimpanzees (Table 2; Supplementary Figure 3) and IFN-γ became significantly associated with infection status with lower levels in infected individuals (P-value =0.019; Table 2; Supplementary Figure 4).

**Table 1.** Mean response values and *P*-values for 15 blood parameters and 8 cytokine/chemokine markers in *Plasmodium*-infected and non-infected chimpanzees. *P*-values were generated from linear models comparing both groups. Statistically significant differences (*P* < 0.05) are marked with an asterisk. White blood cell, platelet, lymphocyte, and monocyte counts are expressed in ×10³/mm³; red blood cell counts in ×10⁶/mm³; hemoglobin in g/dL; hematocrit in %; and neutrophils in ×10⁹/L. GGT, ASAT, and ALAT are expressed in U/L; creatinine in μmol/L; and urea, triglycerides, and cholesterol in mmol/L. Cytokine and chemokine levels were measured in plasma using a Luminex assay and are reported as median fluorescence intensity (MFI).

| Parameter | Mean<br>Non Infected | Mean<br>Infected | P-value |
| --- | --- | --- | --- |
| White blood cells | 9.29 | 9.78 | 0.773 |
| Red blood cells | 4.53 | 4.98 | 0.052 |
| Platelets | 253.79 | 268.54 | 0.688 |
| Neutrophyls | 6.12 | 6.31 | 0.908 |
| Monocytes | 0.48 | 0.49 | 0.914 |
| Lymphocytes | 2.21 | 2.49 | 0.303 |
| Hemoglobine | 12.81 | 13.15 | 0.554 |
| Hematocyte | 36.52 | 38.72 | 0.173 |
| GGT | 19.20 | 13.67 | 0.148 |
| ASAT | 19.04 | 26.09 | 0.145 |
| ALAT | 16.98 | 19.07 | 0.520 |
| Urea* | 0.80 | 2.17 | 0.012 |
| Creatinine* | 86.11 | 67.63 | 0.014 |
| Triglycerides | 1.20 | 1.32 | 0.678 |
| Cholesterol | 4.03 | 3.76 | 0.368 |
| IL4 | 7.18 | 7.58 | 0.495 |
| CCL3* | 3.00 | 21.42 | 0.001 |
| CCL5 | 3223.64 | 4613.58 | 0.065 |
| TNF* | 1.54 | 2.85 | 0.001 |
| IL6 | 1.57 | 2.62 | 0.237 |
| IFNG | 1.36 | 1.23 | 0.599 |
| IL1B* | 0.71 | 1.35 | 0.003 |
| IL10* | 1.89 | 13.73 | 0.002 |

**Figure 3.**
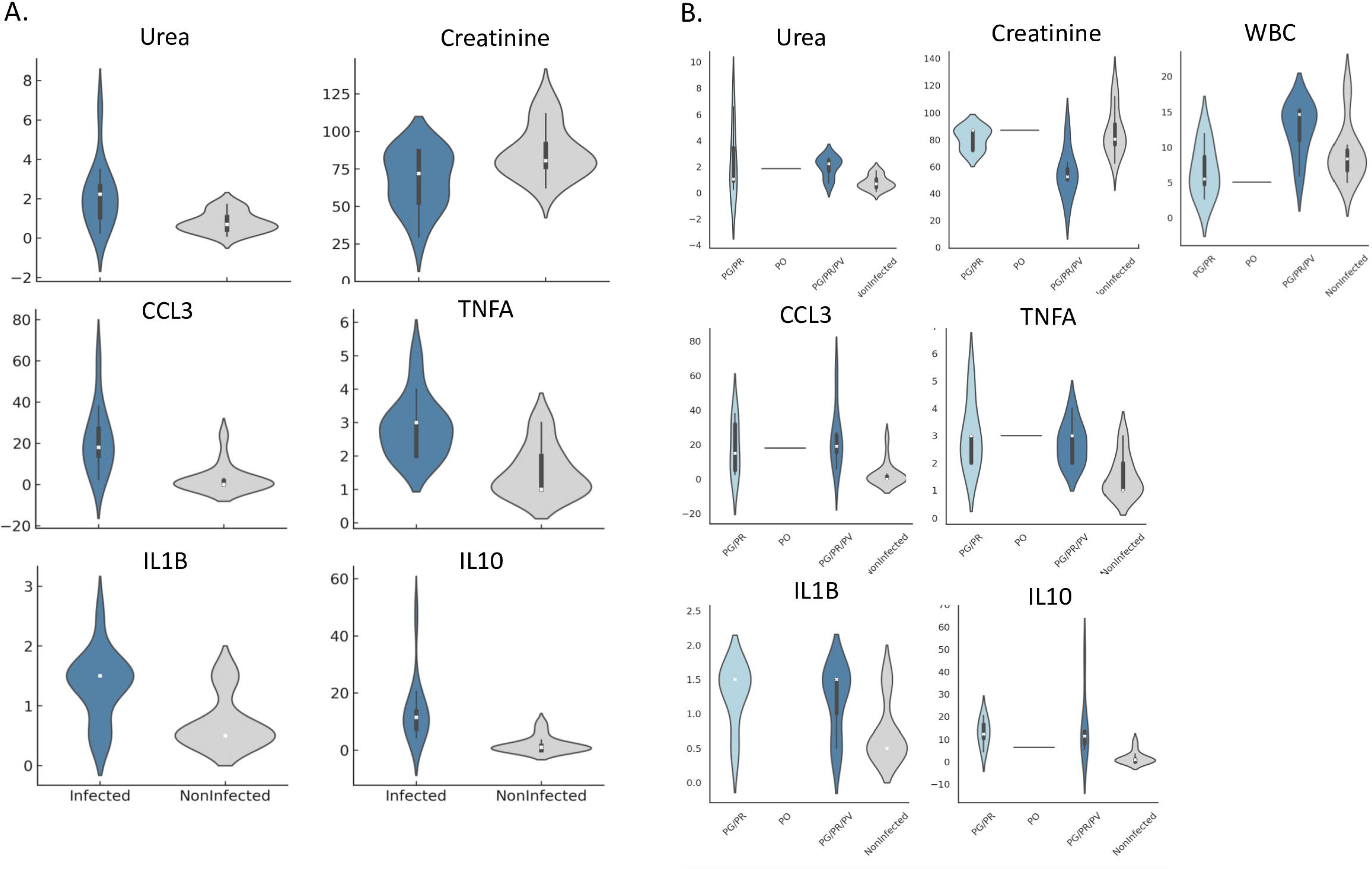
Violin plots of hematological, biochemical and chemokine/cytokine parameters associated significantly with *Plasmodium* infection status in chimpanzees. Each violin plot displays the distribution density for each infection group, with embedded boxplots showing the interquartile range (IQR), central white dots indicating the median, and vertical lines representing the data range (excluding outliers). (A) Distribution of the six parameters that differed significantly between *Plasmodium*-infected and non-infected chimpanzees: urea, creatinine, CCL3, TNF, IL-1β, and IL-10. (B) Distribution of these same parameters across infection types: PG/PR (*P. gaboni* and *P. reichenowi*), PG/PR/PV (triple infection with *P. gaboni*, *P. reichenowi*, and *P. vivax*-like), PO (*P. ovale*-like), and non-infected animals. White blood cell count (WBC) and IFN-γ, which were significantly associated with infection status only when stratified by species group, are also shown. Urea levels (mmol/L) reflect nitrogen waste excretion efficiency; creatinine (μmol/L) serves as a marker of kidney function; WBC counts are expressed in ×10³/mm³. Cytokine and chemokine concentrations (IL-10, TNF, IL-1β, CCL3 and IFN-γ) were measured in plasma using a Luminex multiplex assay and are reported as median fluorescence intensity (MFI), representing relative abundance. Overall, infected chimpanzees—particularly those with multi-species infections—exhibited elevated urea and inflammatory cytokine levels, along with reduced creatinine and WBC counts. Values are normalized residuals; negatives indicate below-average levels

**Table 2.** Mean response values and *P*-values for 15 blood parameters and 8 cytokine/chemokine markers in non-infected chimpanzees and those infected with different *Plasmodium* species. *Plasmodium* species combinations: PG/PR/PV (*P. gaboni*, *P. reichenowi*, *P. vivax*-like) and PG/PR (*P. gaboni*, *P. reichenowi*). *P*-values were obtained from linear models comparing each infected group to the non-infected group. Statistically significant differences (*P* < 0.05) are indicated with an asterisk. White blood cell, platelet, lymphocyte, and monocyte counts are expressed in ×10³/mm³; red blood cell counts in ×10⁶/mm³; hemoglobin in g/dL; hematocrit in %; and neutrophils in ×10⁹/L. GGT, ASAT, and ALAT values are in U/L; creatinine in μmol/L; and urea, triglycerides, and cholesterol in mmol/L. Cytokine and chemokine responses were measured in plasma using a Luminex assay and are reported as median fluorescence intensity (MFI).

| Parameter | Mean Non Infected | Mean PG/PR | Mean PG/PR/PV | P-value |
| --- | --- | --- | --- | --- |
| White blood cells* | 9.29 | 6.72 | 12.64 | 0.045 |
| Red blood cells | 4.53 | 4.75 | 5.09 | 0.120 |
| Platelets | 253.79 | 233.40 | 309.57 | 0.2967 |
| Neutrophils | 6.12 | 3.23 | 8.88 | 0.056 |
| Monocytes | 0.48 | 0.46 | 0.56 | 0.783 |
| Lymphocytes | 2.21 | 2.50 | 2.68 | 0.259 |
| Hemoglobine | 12.81 | 12.44 | 13.41 | 0.503 |
| Hematocryte | 36.52 | 37.00 | 39.01 | 0.396 |
| GGT | 19.20 | 14.4 | 12.41 | 0.323 |
| ASAT | 19.04 | 22.84 | 22.31 | 0.554 |
| ALAT | 16.98 | 14.22 | 18.59 | 0.440 |
| Urea* | 0.80 | 2.46 | 2.00 | 0.025 |
| Creatinine* | 86.11 | 81.04 | 55.28 | 0.001 |
| Triglycerides | 1.20 | 1.79 | 0.98 | 0.117 |
| Cholesterol | 4.03 | 3.72 | 3.67 | 0.549 |
| IL4 | 7.18 | 7.00 | 7.50 | 0.804 |
| CCL3* | 3.00 | 18.50 | 24.00 | 0.003 |
| CCL5 | 3223.64 | 3356.70 | 5098.00 | 0.003 |
| TNF* | 1.54 | 3.00 | 2.71 | 0.382 |
| IL6 | 1.57 | 3.20 | 1.86 | 0.373 |
| IFNG* | 1.36 | 1.00 | 1.14 | 0.019 |
| IL1B* | 0.71 | 1.30 | 1.21 | 0.003 |
| IL10* | 1.89 | 12.90 | 15.36 | 0.003 |

To test whether physiological biomarkers and cytokines predict *Plasmodium* infection status, and to quantify their joint contribution, a ridge-penalized logistic regression was fitted. The model showed balanced discrimination (AUC = 0.80), correctly classifying most infected and non-infected individuals from the combined biomarker and cytokine profiles (Figs. 4A–B). Consistently, the LASSO regression retained five of these six parameters as the strongest predictors of infection status: urea (coefficient = 0.41), IL-10 (0.53), TNF (0.49), CCL3 (0.38), and IL-1β (0.33) (Figure 4C). All parameters were positively associated with infection status. Although creatinine and WBC were not retained in the penalized model, their significance in univariate and species-stratified analyses suggests broader physiological relevance in the host response to *Plasmodium* infection.

**Figure 4.**
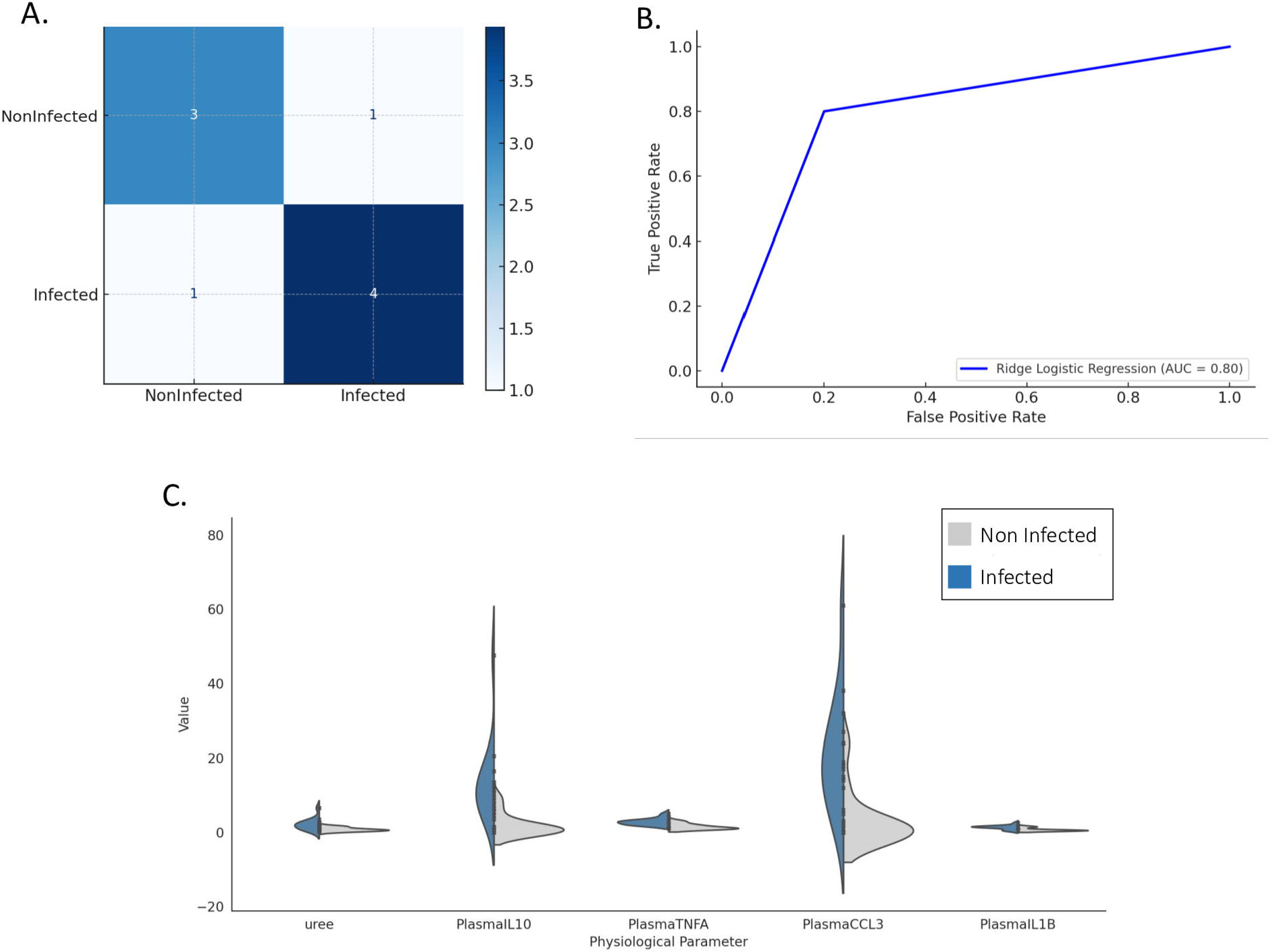
Ridge logistic regression model for predicting *Plasmodium* infection in chimpanzees. (A) Confusion matrix from the ridge logistic regression model predicting infection status based on selected physiological and cytokine parameters. The model correctly classified 3 of 4 non-infected and 4 of 5 infected animals, with one misclassification in each group, indicating balanced predictive performance. (B) Receiver Operating Characteristic (ROC) curve for the same model, based on principal components derived from physiological and cytokine data. The area under the curve (AUC) is 0.80, reflecting good discriminatory ability. (C) Violin plots showing the five parameters retained as significant predictors by LASSO regression: urea, IL-10, TNF, CCL3, and IL-1β. Urea values are expressed in mmol/L; cytokine and chemokine concentrations (IL-10, TNF, CCL3, IL-1β) were measured in plasma using a Luminex assay and are reported as median fluorescence intensity (MFI). Each violin represents the distribution of values by infection status, with internal boxplots indicating the interquartile range (IQR), central white dots showing the median, and vertical lines representing the data range (excluding outliers). These five parameters were consistently identified as the most informative for distinguishing infected from non-infected chimpanzees.

### *Ex vivo* cytokine responses in infected chimpanzees are predominantly anti-inflammatory

Upon *ex vivo* stimulation with PMA or ionomycin directly performed after isolation in the field laboratory, PBMCs isolated from infected chimpanzees produced significantly higher levels of the anti-inflammatory cytokine IL-10 compared to non-infected controls (P-value < 0.05 in both cases; Table 3). This trend remained consistent when analyzing different *Plasmodium* species independently (Table 4).

**Table 3.**
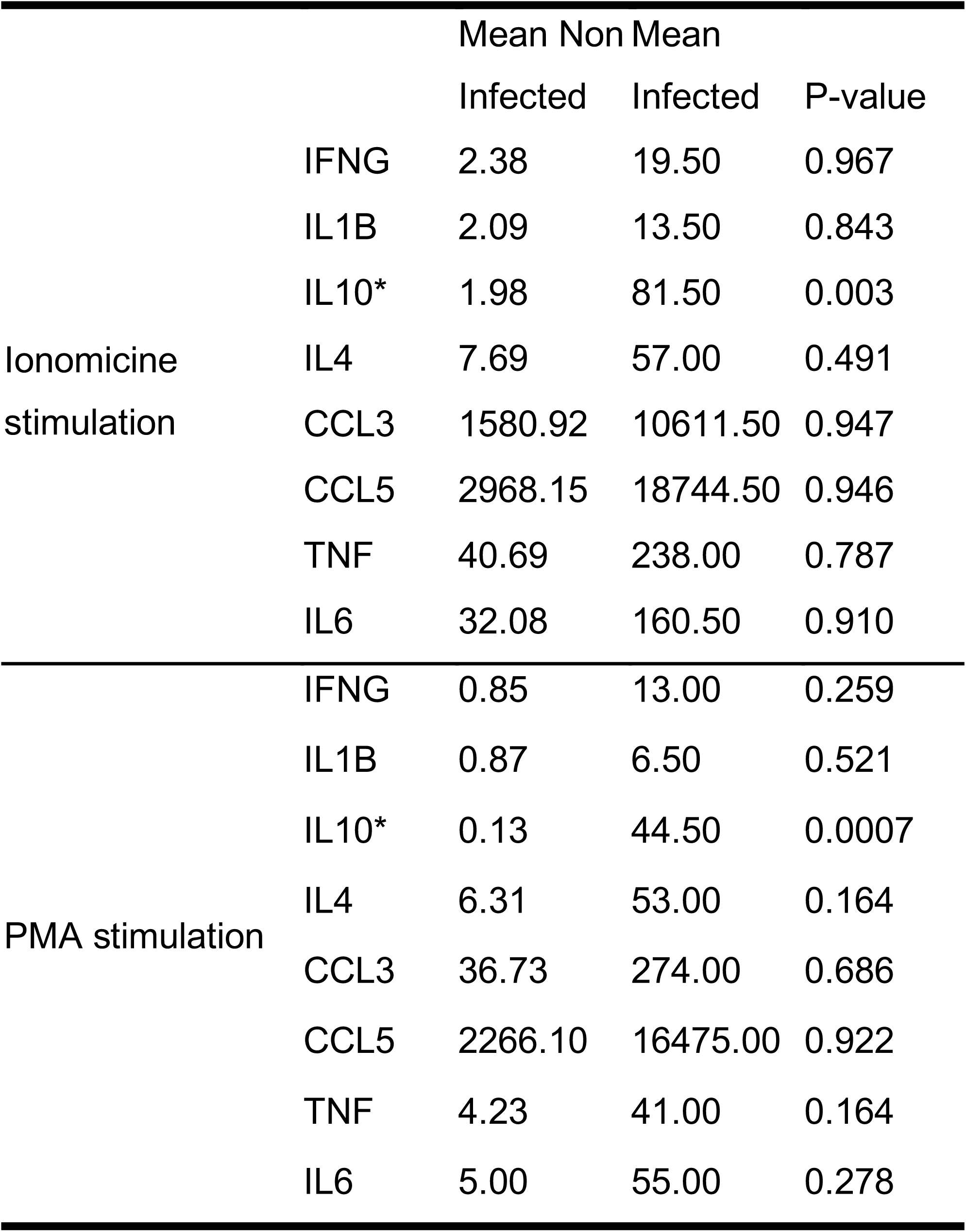
Mean response values and *P*-values for eight cytokine parameters following ionomycin and PMA stimulation in *Plasmodium*-infected and non-infected chimpanzees. *P*-values were derived from linear models comparing cytokine levels between the two groups. Statistically significant differences (*P* < 0.05) are indicated in bold and marked with an asterisk.

**Table 4.** Mean repsonse values and *P*-values for eight cytokine parameters following ionomycin and PMA stimulation in non-infected chimpanzees and those infected with different *Plasmodium* species. *Plasmodium* species combinations: PG/PR/PV (*P. gaboni*, *P. reichenowi*, *P. vivax*-like) and PG/PR (*P. gaboni*, *P. reichenowi*). *P*-values were calculated using linear models comparing each infected group to the non-infected group. Statistically significant differences (*P* < 0.05) are shown in bold and marked with an asterisk.

|  |  | Mean Non | Mean | Mean |  |
| --- | --- | --- | --- | --- | --- |
|  |  | Infected | PG/PR | PG/PR/PV | P-value |
| Ionomicine<br>stimulation | IFNG | 2.38 | 2.67 | 2.67 | 0.967 |
|  | IL1B | 2.09 | 1.83 | 1.83 | 0.843 |
|  | IL10* | 1.98 | 17.83 | 17.83 | 0.003 |
|  | IL4 | 7.69 | 8.00 | 8.00 | 0.491 |
|  | CCL3 | 1580.92 | 1686.80 | 1686.80 | 0.947 |
|  | CCL5 | 2968.15 | 2514.70 | 2514.70 | 0.946 |
|  | TNF | 40.69 | 37.67 | 37.67 | 0.787 |
|  | IL6 | 32.08 | 10.67 | 46.25 | 0.910 |
| PMA stimulation | IFNG | 0.85 | 1.67 | 1.50 | 0.259 |
|  | IL1B | 0.87 | 0.83 | 2.00 | 0.521 |
|  | IL10* | 0.13 | 11.17 | 6.00 | 0.0007 |
|  | IL4 | 6.31 | 7.33 | 7.50 | 0.164 |
|  | CCL3 | 36.73 | 28.67 | 85.00 | 0.686 |
|  | CCL5 | 2266.10 | 2615.5 | 2970.80 | 0.9222 |
|  | TNF | 4.23 | 5.67 | 8.00 | 0.164 |
|  | IL6 | 5.00 | 5.68 | 6.00 | 0.278 |

## Discussion

Although the health effects of *P. falciparum* and *P. vivax* infections in humans are well characterized, the impact of *Plasmodium* infections on great apes remains largely unknown. To address this, we assessed hematological, biochemical and immunological markers in 27 semi-wild chimpanzees in Gabon through field-based sample processing. Obtained results showed that *Plasmodium* infection was highly prevalent (48.15%) among chimpanzees, with younger individuals significantly more infected and those with multi-species infections, particularly PG/PR/PV tri-infections, exhibiting significantly higher parasitemia levels, suggesting both age-dependent susceptibility and additive effects of co-infection (Figure 1). Infected chimpanzees exhibited altered urea and creatinine levels, species-specific changes in white blood cell counts, and elevated concentrations of certain pro-and anti-inflammatory cytokines, indicating systemic immune activation and potential long-term immunomodulation.

Regarding age pattern and infection burden, in our cross-sectional sample, infections were significantly higher in younger chimpanzees, while no older adults tested positive (Figs. 1B, 1D) [24,27–30]. This contrasts with human malaria, where exposure typically yields non-sterilizing (disease-mitigating) immunity; if sustained in larger samples, the chimpanzee pattern is compatible with near-sterilizing immunity accumulating with age/exposure, though alternative explanations (sampling, detectability at low parasitemia, exposure differences) remain possible. A similar age-related decline in *Plasmodium* infection prevalence was reported by De Nys et al. in wild chimpanzees, and by Charpentier et al. in mandrills in Gabon, both suggesting that repeated exposure may lead to partial or effective immunity in African primates [7,31]. Parasitemia increased with infection complexity, being highest in triple-species infections and lowest in *P. ovale-like* mono-infection (Fig. 1C). Consistently with human data, mixed-species infections are often associated with higher parasite densities and more severe outcomes, including anemia and respiratory complications [32–35]. Although we detected no overt clinical signs in this cohort, elevated parasitemia in younger individuals with multi-species infections, particularly those involving *P. vivax-like*, raises the possibility of subclinical impacts, underscoring the need for longitudinal surveillance and immunological profiling in wild ape populations.

Regarding hematological, biochemical and immunological consequences of chronic *Plasmodium* infection in chimpanzees, 23 key markers were compared between infected and uninfected animals, revealing distinct patterns with important implications for disease tolerance and host-pathogen dynamics.

Biochemical analysis revealed that infected animals had an elevated urea-to-creatinine ratio, indicative of pre-renal dysfunction commonly associated with impaired renal hyhpoperfusion in humans. In the context of chronic malaria, kidney function can be significantly affected, potentially leading to complications such as malarial nephropathy and acute kidney injury (AKI) [36]. These renal impairments may result from a combination of hemodynamic disturbances, immune-mediated damage, and systemic effects of prolonged parasitemia. While AKI is more frequently reported in severe malaria cases, emerging evidence suggests that chronic infections can also lead to subclinical or progressive renal dysfunction [36]. In this study, these biochemical and immunological signatures in asymptomatically infected chimpanzees may therefore reflect early or mild forms of renal stress linked to chronic immune activation and parasite persistence.

Regarding immune responses, infected chimpanzees exhibited significantly elevated concentrations of TNF and IL-10 compared to non-infected animals, consistent with reports in humans with asymptomatic *P. falciparum* infections [37]. TNF, a pro-inflammatory cytokine produced primarily by γδ T cells and CD14⁺ monocytes during malaria infection in humans, plays a key role in controlling parasitemia, but can cause damage if not tightly regulated [38–40]. Conversely, IL-10 (anti-inflammatory), primarily produced by IFN-γ⁺ Th1 cells and regulatory T cells in response to chronic antigenic stimulation, counteracts inflammation and limits immunopathology, a phenomenon frequently observed in asymptomatic falciparum malaria [41–44]. Longitudinal monitoring of these chimpanzees over the past 6 years revealed that infected individuals often experienced persistent or recurrent infections (unpublished data), supporting a role for IL-10 in the ongoing regulation of immune responses during chronic exposure. This is consistent with observations in humans living in high-transmission settings, where IL-10 produced by IFN-γ–producing CD4⁺ Th1 cells helps control both immunopathology and clinical symptoms [43,44]. Taken together, these results suggest the existence of a finely regulated balance between pro-inflammatory (TNF) and anti-inflammatory (IL-10) responses in enabling infected chimpanzees to tolerate persistent *Plasmodium* infection while minimizing clinical disease.

Beyond TNF and IL-10, results also showed a significant increase in CCL3 and IL-1β levels among infected chimpanzees. CCL3, a chemokine typically downregulated in asymptomatic malaria in humans [45], has been associated with the development of severe disease, including cerebral and placental malaria [46,47]. Similarly, IL-1β is a hallmark of acute malaria infection and a marker of disease severity in humans [48]. The concurrent upregulation of CCL3 and IL-1β observed in infected chimpanzees may reflect species-specific differences in immune regulation, including leukocyte recruitment (notably neutrophils), inflammatory response thresholds, or parasite clearance mechanisms. These differences could also involve differences in erythrocyte sequestration [49] or parasite-driven immune evasion strategies [4] (Figure 5). The concurrent increase in CCL3 and IL-1β underscores the need to further investigate whether these immune responses confer enhanced parasite control, increased risk of immunopathology, or represent a unique adaptation of chimpanzees to chronic malaria exposure.

**Figure 5.**
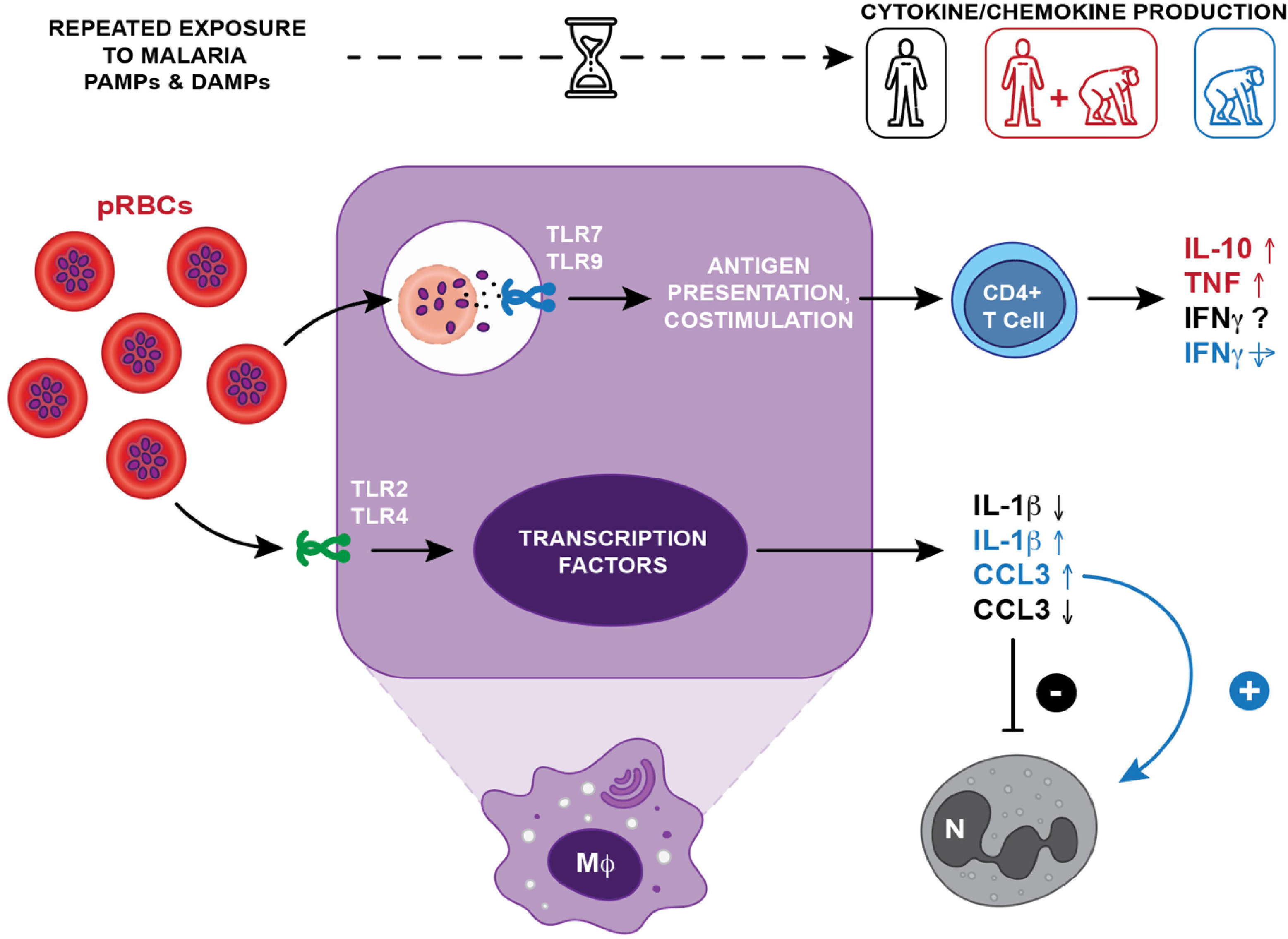
Comparative model of cytokine and chemokine responses in humans and chimpanzees with chronic *Plasmodium* infection. With repeated exposure to *Plasmodium*-parasitised red blood cells (pRBCs), both humans and chimpanzees develop immune tolerance, leading to asymptomatic infections. In humans, an initial pro-inflammatory response (e.g., TNF, IL-1β, CCL3) shifts over time toward a more regulated profile, marked by increased IL-10 production by CD4⁺ T cells and decreased IL-1β and CCL3, leading to an inactivation of neutrophils over time (-). In contrast, IL-1β and CCL3 remain elevated in chronically infected chimpanzees, suggesting sustained immune activation involving neutrophils (+). Effector mechanisms such as phagocytosis, phagolysosome activation, antigen presentation, T cell co-stimulation, and IFN-γ production by CD4⁺ T cells are enhanced in humans through Toll-like receptor (TLR) signaling, which recognizes *Plasmodium* pathogen-associated molecular patterns (PAMPs) and damage-associated molecular patterns (DAMPs) (PMID: 25324127). These mechanisms help maintain parasite control without clinical symptoms. The role of IFN-γ in this regulatory sequence remains debated in humans but similar levels are seen in both healthy and infected chimpanzees and IFN concentrations decrease in bi-and tri-infected animals compared to controls. Cytokine and chemokine profiles observed in our animal cohort are compared to known responses in humans: **black** = human-specific responses; **red** = shared trends in humans and chimpanzees; **blue** = responses observed only in chimpanzees (adapted from PMID: 24743880). Mφ = macrophage; N = neutrophil.

When comparing chimpanzees with bi-and tri-species *Plasmodium* infections to non-infected ones, similar profiles emerged. Tri-infected animals (*P. gaboni*, *P. reichenowi*, and *P. vivax*-like) displayed higher white blood cell counts, higher urea and lower creatinine levels, and a cytokine profile marked by increased TNF, CCL3, IL-1β, and IL-10, relative to bi-infected animals (*P. gaboni* and *P. reichenowi*). Notably, IFN-γ levels were significantly lower in multiple infections, suggesting a dampened Th1 response. These patterns suggest that co-infection with *P. vivax*-like may alter immune regulation, potentially shifting the host response toward a more anti-inflammatory or immunomodulatory profile, possibly through enhanced IL-10-mediated suppression of pro-inflammatory pathways combined with reduced IFN-γ activity. This may reflect immune exhaustion, tolerance, or parasite-driven immune evasion mechanisms in the context of chronic, multi-species malaria infections. Overall, these results highlight the possible role of *P. vivax-like* co-infection in further modulating immune responses, possibly enhancing immune tolerance and facilitating persistent, asymptomatic multi-species malaria infection in chimpanzees.

PBMCs from infected chimpanzees exhibited significantly higher IL-10 upon stimulation with ionomycin or PMA compared to non-infected controls, while pro-inflammatory cytokine production (TNF, IFN-γ, IL-6) remained low. This result further supports our earlier observations of elevated systemic IL-10 levels in infected animals, underscoring the prominence of an immunoregulatory state in response to chronic *Plasmodium* infection. Such a profile is consistent with reports from *P. falciparum*-endemic regions, where repeated infections dampen monocyte activation and inflammatory responses [50,51]. Such IL-10–dominated profiles may reflect immune tolerance or epigenetic reprogramming of monocytes, as seen in asymptomatic malaria cases in Malian adults and children, potentially increasing susceptibility to secondary infections [45]. Together, these findings highlight the central role of IL-10 in orchestrating both systemic and cellular immune regulation during chronic malaria exposure in chimpanzees. Serological data on past exposures would help clarify the role of chronic infection in shaping this regulatory phenotype.

This study advances our understanding of the hematological and immune landscape in wild chimpanzees with asymptomatic *Plasmodium* infections, revealing striking similarities to the immune responses observed in human malaria. Contrary to the notion that chronic malaria is immunologically silent, our results suggest that infected chimpanzees undergo significant shifts in cytokine profiles, particularly elevated systemic and cellular IL-10 levels. This pattern suggests a state of immunosuppressive priming, which may enable the host to tolerate persistent parasitemia, but could also compromise defenses against other pathogens. In addition, the detection of subtle alterations in renal function, especially in chimpanzees with tri-species infections, further indicates that even asymptomatic malaria can exert physiological impacts.

While all confirmed infections in our cohort were asymptomatic, regardless of parasitemia, disease expression may vary depending on the host and parasite species. Clinical cases of *P. reichenowi* in chimpanzees [12], and of *Plasmodium pitheci* in rehabilitated Bornean orangutans have been documented [11]. Persistent immune stimulation may lead to immune exhaustion or paralysis, potentially increasing susceptibility to other infections. This highlights the importance of ongoing immune monitoring, particularly in juveniles with multi-species infections. Accordingly, management plans should consider immune health alongside population size, as asymptomatic infections can carry hidden costs. Field-feasible panels, complete blood counts and simple immune assays (e.g., inflammatory markers or parasite-specific antibodies), could serve as indicators of population health, especially in juveniles with multi-species infections.

Finally, it has to be added, that we acknowledge limitations in this study, particularly the modest sample size, restriction to asymptomatic individuals, and a limited bio-markers panel. To overcome such limit, it would be interesting in the future to expand immune profiling to include additional regulatory markers (e.g., IL-12, TGF-β) and implementing longitudinal and serological monitoring for a more comprehensive understanding of immune memory, reinfection, and long-term health outcomes in these endangered populations.

In summary, this study suggests that ostensibly asymptomatic malaria in chimpanzees is not benign: immune modulation varies with infection complexity and age, a pattern compatible with virulence expressed as subclinical costs (e.g., anemia risk, altered inflammatory tone, reduced condition). Together with accumulating reports in other primates that malaria affects physiology and performance even without overt illness [52], this argues for integrating longitudinal health surveillance into conservation practice.

## Methods

### Ethical statement

All animal procedures were conducted in strict accordance with national and international guidelines for animal welfare to ensure the well-being of the animals. Veterinary staff from La Lékédi Park and the Centre Interdisciplinaire de Recherches Médicales de Franceville (CIRMF) supervised all health assessments and blood collection. Ethical approval was obtained from the government of the Republic of Gabon and the Animal Life Administration of Libreville (CITES 00956), as well as the Animal Welfare and Ethical Review Body of the London School of Hygiene & Tropical Medicine (LSHTM; 0020/2013/SG/CNE).

### Study sites and sample collection

Blood samples were collected from 27 chimpanzees as part of routine veterinary health assessments at two sites in Gabon: La Lékédi Park (N=9) and the Primatology Center at CIRMF, Franceville (N=18) (Figure 1A). Sampling took place during peak malaria transmission seasons—October 2018 at La Lékédi (5 females, 4 males) and May 2019 at CIRMF (11 females, 11 males). Four animals were excluded due to incomplete data, yielding a final sample size of 27. During assessments, chimpanzees were anesthetized with an intramuscular combination of medetomidine (0.03–0.06 mg/kg IM) and ketamine (2–6 mg/kg IM; Imalgene®) delivered via blowpipe; for adults, ketamine was typically 3 mg/kg within this range, with medetomidine adjusted according to clinical judgement. Blood was collected from the iliac vein within 15 minutes of sedation to minimize stress-related alterations in hematological parameters. Three blood samples were obtained from each animal using dry, EDTA, and heparin tubes for subsequent analysis. Age, weight, and rectal temperature were also recorded.

### *Laverania* species diagnosis

To reduce host DNA within samples, 4 mL of EDTA-collected chimpanzee blood was diluted with PBS and passed through CF11 cellulose columns for leukocyte depletion [53]. Genomic DNA was extracted using the DNeasy Blood and Tissue Kit (Qiagen, France) following the manufacturer’s instructions. *Plasmodium* species were identified by nested PCR amplification and sequencing of the cytochrome *b* (cyt-b) gene using two primer sets: DW2–DW4 [54]. PCR products were visualized via 1.2% agarose gel electrophoresis and quantified with a Qubit fluorometer.

Sequencing was then performed using high-throughput technology (MinION) to allow the identification of mixed-infections. DNA libraries were prepared using the Ligation Sequencing Kit (SQK-NBD114.96) and Native Barcoding Kit 96 V4, then sequenced on a MinION Mk1C device (Oxford Nanopore Technologies) with FLO-MIN114 (R10.4.1) flow cells, targeting 100,000–200,000 reads per sample. For species identification, a standardized random subsample of 10,000 reads per chimpanzee was BLASTed against a custom Cytb database. This sub-sampling approach optimized computational efficiency and ensured cross-sample comparability without compromising species-level resolution.

### Plasmodium vivax-like diagnostic

Due to the initial PCR protocol’s limited sensitivity for detecting *P. vivax*-like parasites, all DNA samples were additionally screened using a conventional PCR targeting the mitochondrial cytochrome oxidase I (*cox1*) gene, following the protocol of Liu et al. [25]. Amplified products were resolved on 1.5% agarose gels and sequenced by Eurofins MWG. Resulting sequences were analyzed using BLAST for species identification (https://blast.ncbi.nlm.nih.gov/Blast.cgi).

### *Plasmodium* parasitemia quantification

Parasite quantification was performed by quantitative PCR (qPCR), comparing sample cycle threshold (Ct) values to a standard curve derived from *P. falciparum* ring-stage parasites. Control parasitemia was determined via Giemsa-stained smear counts across eight distant fields, each containing ∼200 red blood cells (RBCs), using the following formula:

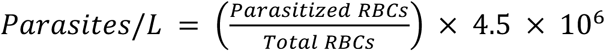

This formula is the standard conversion from % parasitaemia on a thin film to parasites per volume of blood. Parasite density on thin films was computed as parasites/µL = (% parasitized RBCs) × (RBC count per µL), using an assumed RBC count of 4.5×10⁶/µL when an individual CBC was unavailable (CDC DPDx; BSH guideline). Values per liter were obtained by multiplying parasites/µL by 10⁶.

Genomic DNA from standards and samples was extracted using the DNeasy Blood and Tissue Kit (Qiagen, France) following the manufacturer’s protocol. Parasitemia was then measured using a custom, unpublished qPCR assay targeting the *Plasmodium* cytochrome *b* gene, allowing the amplification also of *P. vivax-like*, with primers ParaF (5′-CTATGCTTTATTATGGATTGGATG-3′) and ParaR (5′-GAGCTGTAATCATAATGTGTTC-3′). Reactions (20 μL) contained one μL of DNA, 10 μM of each primer, and 1× Evagreen. Amplification was performed on a LightCycler 96 with the following cycling conditions: 95 °C for 10 min; 40 cycles of 95 °C for 15 s, 60 °C for 20 s, 72 °C for 20 s; followed by 95 °C for 10 s and 55 °C for 1 min. Standards were run in duplicate across a dilution range from 300,000 to 1 parasite/μL. Sample parasitemia was estimated by extrapolating Ct values against the standard curve using LightCycler 96 software.

### Chimpanzees blood group status

Given the known association between ABO blood group and malaria susceptibility in humans [55], ABO blood group and rhesus (Rh) status were determined for each chimpanzee using the SERAFOL® ABO+D kit within 3 hours of blood collection. Blood samples were tested for agglutination with specific antisera to identify blood group and Rh status.

### Blood numeration and serum biochemical analyses

To evaluate the impact of *Plasmodium* infection on chimpanzee health, we assessed eight hematological parameters commonly altered in human malaria, including total white blood cell (WBC), red blood cell (RBC), platelet, neutrophil, monocyte, and lymphocyte counts, as well as hemoglobin and hematocrit levels, using an automated hematology analyzer (Hematology Analyser ACT 10, Beckman Coulter, USA). Complete blood counts were performed within 90 minutes of sample collection at the CIRMF Medical Analysis Center. Renal (urea, plasma creatinine) and hepatic (ALAT, ASAT, GGT) function markers were also measured, given the known impact of malaria on these organs in humans, using a fully automated clinical chemistry analyzer (Hitachi Model 902 Automatic Analyser, Roche Diagnostics, France), following the manufacturer’s protocols. In addition, total cholesterol and triglyceride levels were analyzed, given the auxotrophic dependence of *Plasmodium* parasites on host cholesterol.

### Peripheral blood mononuclear cell (PBMC) stimulation

To assess immune status, we used phorbol myristate acetate (PMA) and ionomycin stimulation, a widely established method for inducing immune responses in vitro across multiple species, including humans [56]. This approach triggers rapid secretion of proinflammatory cytokines and chemokines within six hours of stimulation [57]. For each animal, 4 mL of heparinized whole blood was processed within 3 hours of collection. Samples were centrifuged (800 × g, 20 min), and plasma was aliquoted and stored at –80 °C. The buffy coat, containing peripheral blood mononuclear cells (PBMCs), was transferred to 24-well plates and stimulated with PMA (25 ng/mL) and ionomycin (1 μg/mL) for 6 hours. After incubation, the cells were centrifuged, and the supernatants were collected and stored at –80 °C for subsequent analysis of cytokines and chemokines.

### Plasma level assessment of cytokines and chemokines

To investigate immune responses associated with malaria infections, we selected eight cytokines and chemokines known to be involved in malaria-associated inflammation and pathogenesis in humans, as well as immunomodulation in asymptomatic infections [58,59]. The selected markers—TNF, IL-6, IFN-γ, IL-1β, IL-4, CCL3/MIP-1α, CCL5/RANTES, and IL-10—were quantified using the Cytokine & Chemokine 30-Plex NHP ProcartaPlex™ Panel (Invitrogen) in plasma samples, including those from stimulated PBMCs. Plasma samples were thawed before cytokine and chemokine levels were quantified using a commercially available multiplex bead assay kit (Primate Magnetic Luminex Assay, Biotechne, Minneapolis, USA) and quantitative suspension array technology (Luminex Corp, Austin, USA). The assay was performed according to the manufacturer’s protocols by individuals who were blinded to the study endpoints. Median fluorescence intensity (MFI) values of samples achieving a bead count of at least 50 per analyte were acquired and background-adjusted using the Luminex MAGPIX© system and xPONENT 4.2 software.

### Statistical analyses

All statistical analysis have been performed using R4.3.1 software [60].

#### (a) Data preparation and quality control

The dataset comprised 15 hematological and 8 cytokine/chemokine parameters from chimpanzees sampled in Gabon, along with demographic variables (age, weight, sex, temperature, blood group) and *Plasmodium* infection status. Before analysis, continuous variables were screened for missing values and outliers, and standardized using z-scores where appropriate for multivariate analyses using the tidyverse R package [61].

#### (b) Global comparison of infected vs. non-infected animals

To assess physiological differences associated with *Plasmodium* infection, all infected animals were compared as a single group to non-infected chimpanzees. Mean values were calculated for each of the 23 physiological parameters (15 hematological parameters, 8 cytokines and chemokines), and Welch’s t-tests were used to compare group means, accounting for unequal variances and sample sizes. Statistical significance was defined as *p* < 0.05. Associations between demographic variables and infection status were assessed using Welch’s t-tests for continuous variables (age, weight, temperature) and t-tests for categorical variables (sex, blood group). Because infected chimpanzees were, on average, younger than non-infected individuals, we accounted for potential confounding by including age as a covariate in logistic regression models when testing associations between infection status and physiological parameters.

Data distributions and group differences were visualized with the ggplot2 package [62] with violin plots displaying kernel density estimates and internal boxplots (median and IQR), colored by infection status (light grey = non-infected; steel blue = infected). Statistically significant differences were annotated with asterisks (P-value < 0.05). To explore global variation in physiological profiles, Principal Component Analysis (PCA) was conducted separately for blood and cytokine parameters using the prcomp() function on z-score standardized values, with shaded ellipses representing 60% confidence regions from a bivariate normal fit to PC1–PC2 scores per group. Variables with Pearson correlation coefficients > 0.9 were excluded to reduce multicollinearity [63].

#### (c) Subgroup comparison by Plasmodium species combinations

To investigate whether specific *Plasmodium* species or species combinations differentially influenced physiological responses, infected animals were subdivided into two groups: PG/PR, and PG/PR/PV (PO being alone was not considered). Each group was independently compared to the non-infected group using Welch’s t-tests. For each comparison, the direction of change, P-value, and statistical significance (P-value < 0.05) were recorded. In addition, violin plots were used to visualize all four groups and PO, with distinct colors assigned: PG/PR (light blue), PO (medium blue), PG/PR/PV (dark blue), and non-infected (light grey). To explore global variation in physiological profiles, Principal Component Analysis (PCA) was conducted separately for blood and cytokine parameters using the prcomp() function on z-score standardized values, with shaded ellipses representing 60% confidence regions from a bivariate normal fit to PC1–PC2 scores per group. Variables with Pearson correlation coefficients > 0.9 were excluded to reduce multicollinearity.

#### (d) Penalized regression and predictive modeling of infection status

To identify physiological markers predictive of *Plasmodium* infection (all species included) in wild chimpanzees, we applied penalized logistic regression using the LASSO (Least Absolute Shrinkage and Selection Operator) method implemented via the glmnet package in R [64]. All predictor variables, including hematological and cytokine parameters, were standardized to z-scores. Age was included as a covariate to control for potential confounding. The optimal regularization parameter (λ) was selected using 10-fold cross-validation, and variables with non-zero coefficients at λmin were retained as key predictors.

Five variables were identified as most informative: urea, IL-10, TNF, CCL3, and IL-1β. Violin plots were generated using ggplot2 to visualize the distributions of urea and cytokines (IL-10, TNF, CCL3, and IL-1β) by infection status, with urea expressed in mmol/L and cytokines reported as median fluorescence intensity (MFI) from plasma Luminex assays. Each plot shows full value distributions with embedded boxplots (median, IQR, and range excluding outliers). Model performance was evaluated using a confusion matrix and ROC curve (p_ROC package), correctly classifying 80% infected and 75% non-infected animals, with an AUC of 0.80, indicating strong discriminatory power [65].

## Acknowledgements

This study was supported by ANR T-ERC EVAD, ANR JCJC GENAD, and CNRS. SCW is funded by grants from the National Institute of Allergy and Infectious Diseases of the National Institutes of Health (NIH, award numbers U19AI089676 and U19AI181587) and the UK Medical Research Council (award number MR/S009450/1). The content is solely the responsibility of the authors and does not necessarily represent the official views of the NIH.

## Data availability

The sequences of *P. vivax-like* reported in this study were deposited in GenBank under the following accession numbers (PV146376, PV146375, PV146364, PV146345, PV146323, PV146314, PV146309)., The MinION sequencing data generated in this study have been deposited in GenBank under the following accession numbers, as part of BioProject PRJNA1299386. The raw data supporting the conclusions of this article is available upon request.

