## Supplementary File for "Hidden hematological, biochemical and immune costs of asymptomatic malaria infections in semi-wild chimpanzees"

### Supplementary Tables

**Supplementary Table 1. Summary of BLAST results for *Plasmodium* species detection in chimpanzee samples using MinION sequencing.** This table presents the results of a BLAST analysis performed on 10,000 randomly selected reads from MinION sequencing data. Each row corresponds to a different chimpanzee sample, and the columns represent the number of reads matching specific *Plasmodium* species. PCR amplification of the cytochrome B gene was initially used to detect the presence of *Plasmodium* species, followed by MinION sequencing. The species identified include *P. adleri*, *P. gaboni*, *P. ovale-like*, *P. praefalciparum*, *P. reichenowi*, *P. vivax-like*, *P. billcollinsi*, *P. malariae-like*, and *P. blacklocki*. The "Non-identified" column indicates the number of reads that could not be confidently assigned to any specific *Plasmodium* species. The final column totals the number of identified reads for each sample. Names followed by an asterisk (*) indicate individuals PCR-positive for *P. vivax-like*.

| Name | *P. adleri* | *P. gaboni* | *P. ovale-like* | *P. praefaciparum* | *P. reichenowi* | *P. vivax-like* | *P. billcolinsi* | *P. malariae-like* | *P. blackloki* | Non-idenfied | Total |
|---|---|---|---|---|---|---|---|---|---|---|---|
| Tarzan | 2 | **2465** | 2 | 1 | **7518** | 1 | 0 | 1 | 2 | 8 | 9992 |
| Tonic | 3 | **8714** | 201 | 21 | **1034** | 13 | 5 | 5 | 0 | 4 | 9996 |
| Ogooue | 8 | **6158** | 11 | 5 | **3798** | 10 | 1 | 0 | 0 | 9 | 9991 |
| Ebene | 2 | **5871** | 2 | 8 | **4106** | 0 | 0 | 0 | 0 | 11 | 9989 |
| Felix | 5 | **1817** | 6 | 11 | **8141** | 7 | 2 | 6 | 0 | 5 | 9995 |
| Malimbe | 1 | **7869** | 3 | 12 | **2107** | 6 | 0 | 0 | 0 | 2 | 9998 |
| Flore | 1 | **459** | 3 | 2 | **9524** | 4 | 0 | 0 | 0 | 7 | 9993 |
| Cerise | 2 | **8114** | 4 | 2 | **1860** | 9 | 0 | 0 | 0 | 9 | 9991 |
| Nikita | 0 | **2263** | 51 | 3 | **7653** | 6 | 0 | 12 | 1 | 11 | 9989 |
| Charly | 0 | **1759** | 5 | 19 | **8196** | 13 | 0 | 0 | 1 | 7 | 9993 |
| Nzigou | 10 | 22 | **9767** | 1 | 94 | 31 | 0 | 9 | 0 | 66 | 9934 |
| Wonga | 4 | **5221** | 1 | 10 | **4652** | 96 | 7 | 0 | 0 | 9 | 9991 |
| Ebea | 0 | **3603** | 7 | 5 | **6356** | 21 | 0 | 0 | 0 | 8 | 9992 |

**Supplementary Table 2. Impact of demographic and physiological variables on malaria-infection status in chimpanzees in Gabon.** This table presents the results of t-tests conducted to determine if age (in months), weight (in kg), sex, blood group, and temperature (in °C) have a significant impact on the infection status (infected vs. non-infected) in chimpanzees. For each variable, the t-statistic value and the corresponding *P*-value are reported.

| Variable | t-statistic | *P*-value |
| --- | --- | --- |
| Age (months) | -5,01 | 0,000036* |
| Weight (kg) | -1,45 | 0,16 |
| Sex | -0,95 | 0,35 |
| Blood group | -0,22 | 0,82 |
| Temperature (°C) | 1,01 | 0,32 |

### Supplementary Figure legends

**Supplementary Figure 1. Violin plots of blood parameters in *Plasmodium*-infected and non-infected chimpanzees.** This figure displays the distribution of 15 blood parameters measured in *Plasmodium*-infected and non-infected chimpanzees. Each violin plot shows the full distribution of values, with an embedded box plot indicating the interquartile range (IQR), the median (central white dot), and the data range (excluding outliers). ALAT, ASAT, and GGT values are expressed in units per liter (U/L). Cholesterol (Chol), triglycerides (TG), and urea concentrations are reported in mmol/L. Creatinine (creat) is expressed in µmol/L. White blood cells (WBC), platelets (PQT), lymphocytes (Lymph), and monocytes (Mono) are presented in $\times 10^3/mm^3$. Red blood cell count (RBC) is expressed in $\times 10^6/mm^3$. Hemoglobin (Hb) is reported in g/dL, and hematocrit (Ht) in percentage (%). Neutrophils (Neut) are expressed in $\times 10^9/L$. These plots allow visual comparison of central tendencies and distributional differences in blood markers between infected and non-infected animals, highlighting potential physiological changes associated with *Plasmodium* infection. Values are normalized residuals; negatives indicate below-average levels

**Supplementary Figure 2. Violin plots of cytokine and chemokine levels in *Plasmodium*-infected and non-infected chimpanzees.** This figure presents the distribution of eight cytokine and chemokine parameters measured in plasma samples from chimpanzees infected with *Plasmodium* and those that were not. Cytokine and chemokine response levels were assessed using a Luminex multiplex assay, and values are reported as median fluorescence intensity (MFI). Each violin plot displays the full distribution of values for each biomarker, with an internal box plot showing the interquartile range (IQR), median (white dot), and data range (excluding outliers). These plots highlight distributional and central tendency differences in immune profiles between infected and non-infected animals. Values are normalized residuals; negatives indicate below-average levels

**Supplementary Figure 3. Violin plots of blood parameter levels measured in non-infected chimpanzees and in chimpanzees infected with different *Plasmodium* species combinations.** Infection categories include PG/PR/PV (infected with *P. gaboni*, *P. reichenowi*, and *P. vivax*-like), PG/PR (infected with *P. gaboni* and *P. reichenowi*), and PO (infected with *P. ovale*-like). Hematological parameters were measured in blood samples collected from each animal. ALAT, ASAT, and GGT values are expressed in units per liter (U/L). Cholesterol (Chol), triglycerides (TG), and urea concentrations are reported in mmol/L. Creatinine (creat) is expressed in µmol/L. White blood cells (WBC), platelets (PQT), lymphocytes (Lymph), and monocytes (Mono) are presented in $\times 10^3/mm^3$. Red blood cell count (RBC) is expressed in $\times 10^6/mm^3$. Hemoglobin (Hb) is reported in g/dL, and hematocrit (Ht) in percentage (%). Neutrophils (Neut) are expressed in $\times 10^9/L$. Each violin plot illustrates the distribution of values, with internal boxplots representing the median and interquartile range, highlighting differences in blood profiles across infection statuses. Values are normalized residuals; negatives indicate below-average levels

**Supplementary Figure 4. Violin plots of eight cytokine and chemokine parameter levels measured in non-infected chimpanzees and in chimpanzees infected with different *Plasmodium* species combinations.** Infection categories include PG/PR/PV (infected with *P. gaboni*, *P. reichenowi*, and *P. vivax*-like), PG/PR (infected with *P. gaboni* and *P. reichenowi*), and PO (infected with *P. ovale*-like). Cytokine and chemokine response levels were quantified from plasma samples using Luminex assays and are expressed in median fluorescence intensity (MFI). Each violin plot illustrates the distribution of values, with internal boxplots representing the median and interquartile range, highlighting differences in cytokine and chemokine responses across infection groups. Values are normalized residuals; negatives indicate below-average levels.
