## Supplementary figures and images for "Hidden hematological, biochemical and immune costs of asymptomatic malaria infections in semi-wild chimpanzees"

### Supplementary Figure 1

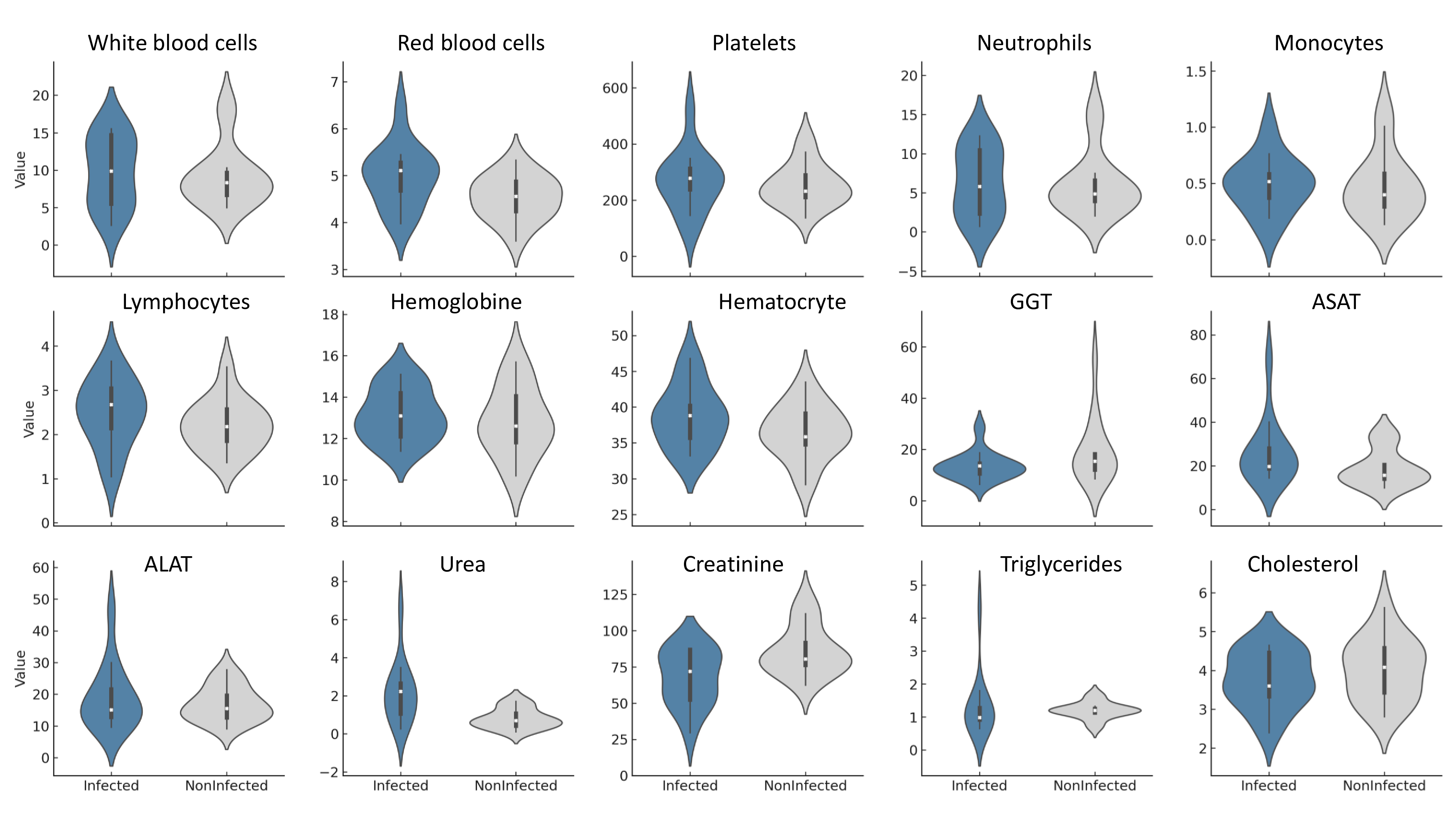

### Supplementary Figure 2

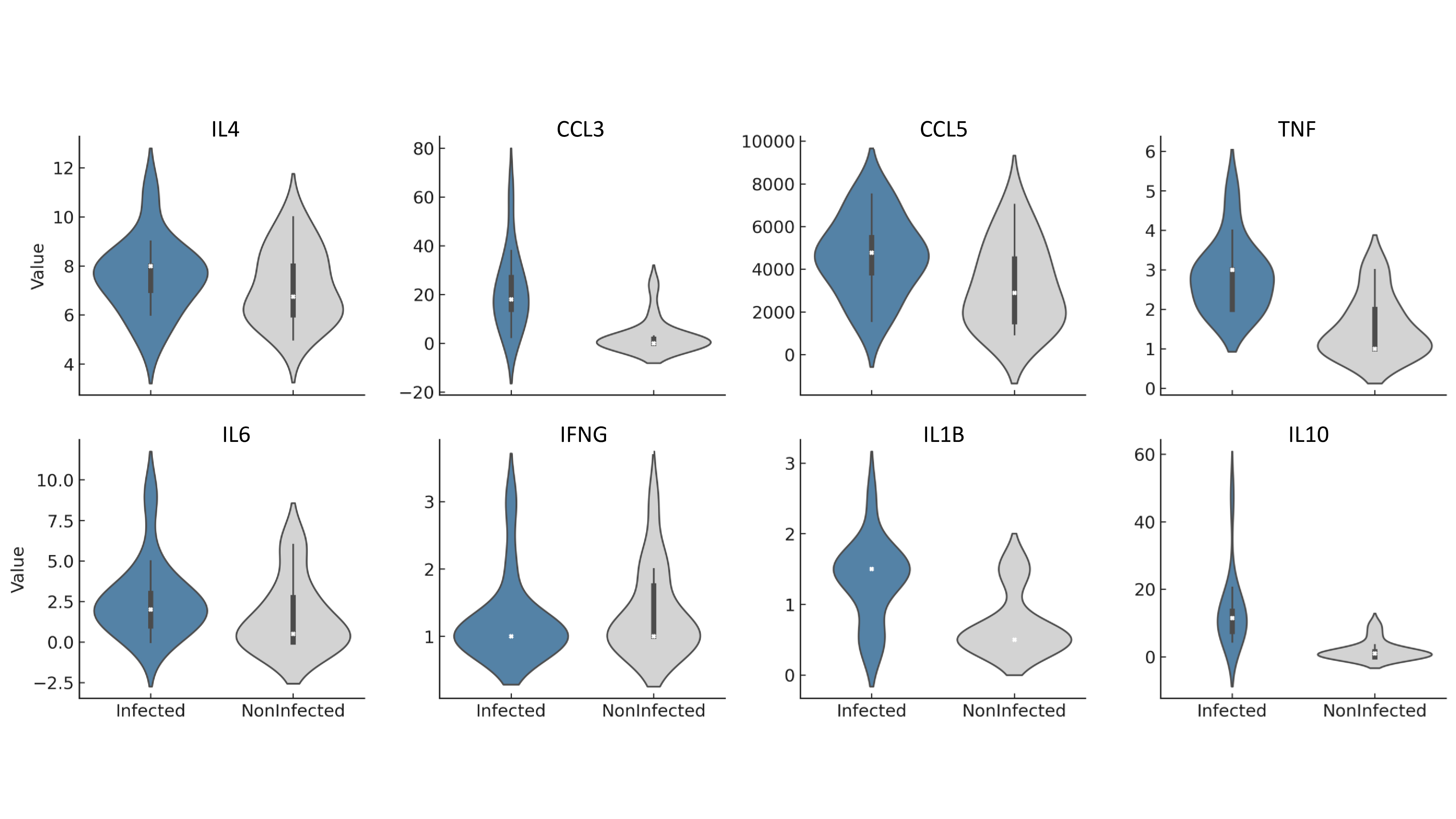

### Supplementary Figure 3

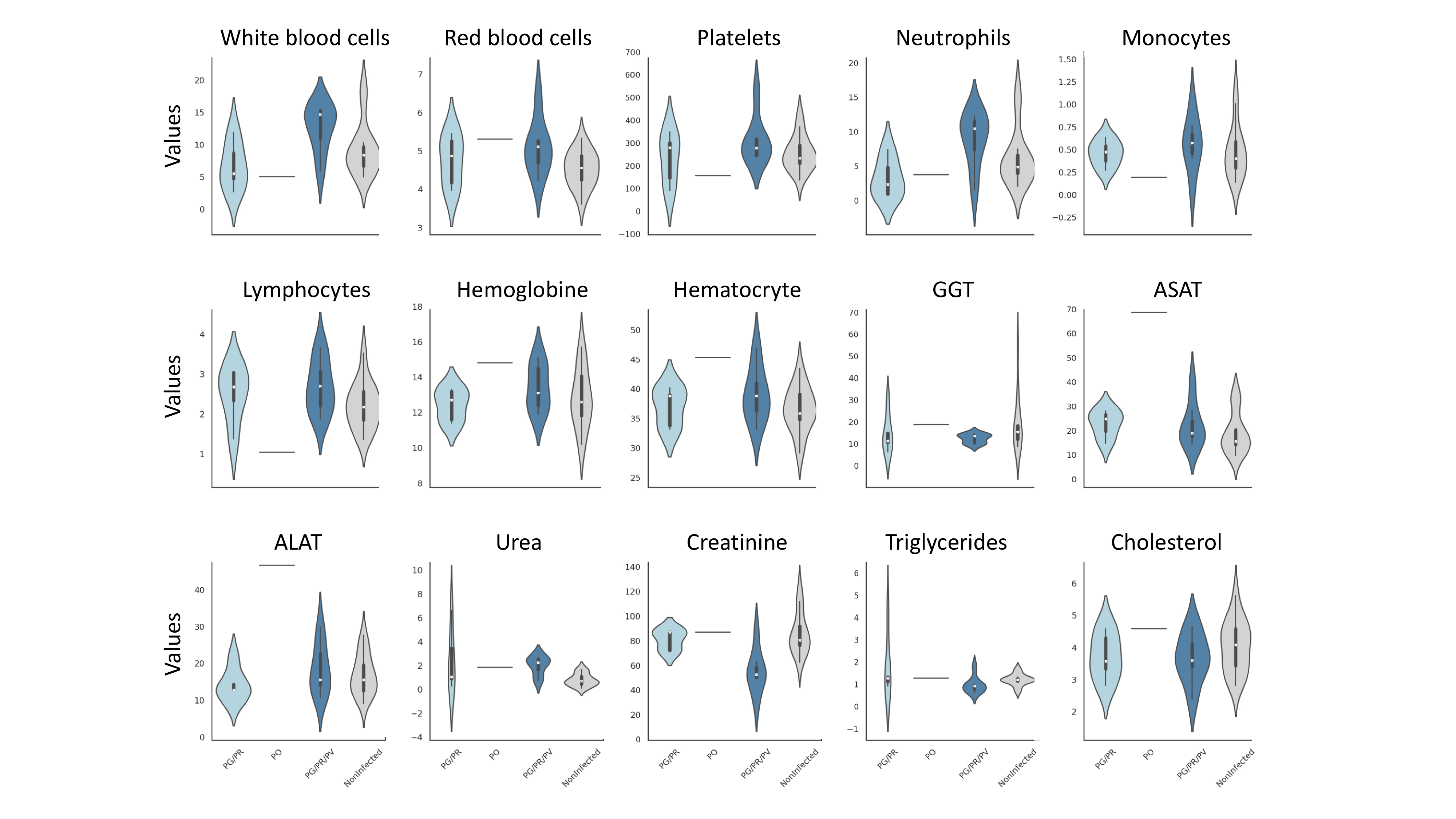

### Supplementary Figure 4

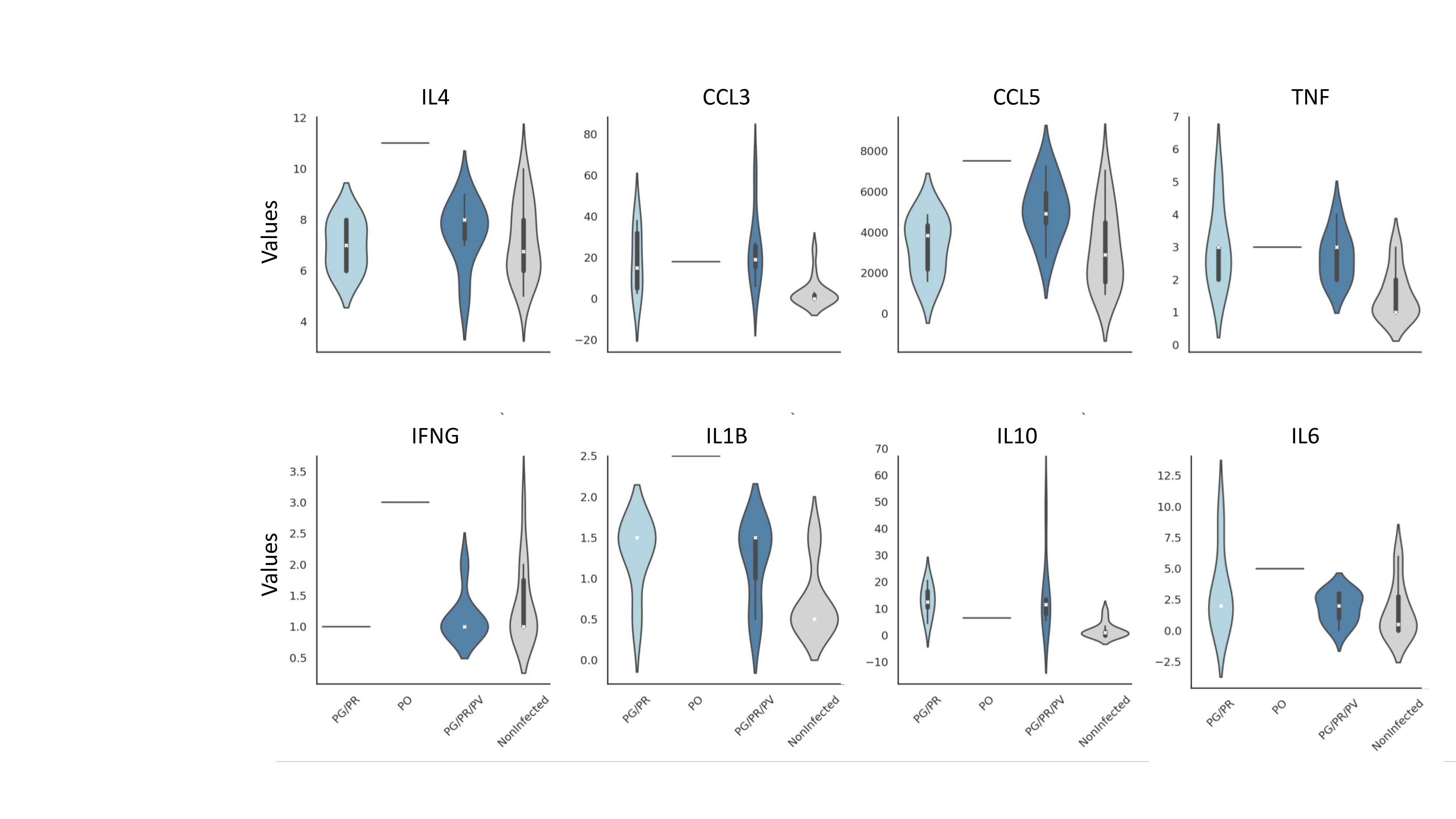
